# Three-dimensional structure of the single domain cupredoxin AcoP

**DOI:** 10.1101/2022.03.21.484586

**Authors:** Magali Roger, Philippe Leone, Ninan Blackburn, Sam Horrell, Tadeo Moreno Chicano, Marie-Thérèse Giudici-Orticoni, Luciano A. Abriata, Greg L. Hura, Michael A. Hough, Giulano Sciara, Marianne Ilbert

## Abstract

Cupredoxins are widely occurring copper-binding proteins with a typical Greek-key beta barrel fold. They are generally described as electron carriers that rely on a T1 copper center coordinated by four ligands provided by the folded polypeptide. The discovery of novel cupredoxins demonstrates the high diversity of this family, with variations in term of copper-binding ligands, copper center geometry, redox potential, as well as biological function. AcoP is a periplasmic protein belonging to the iron respiratory chain of the acidophilic bacterium *Acidithiobacillus ferrooxidans*. AcoP presents original features: highly resistant to acidic pH, it possesses a constrained green-type copper center of high redox potential. To understand the unique properties of AcoP, we undertook structural and biophysical characterization of wild-type AcoP and of two Cu-ligand mutants (H166A and M171A). The crystallographic structure of AcoP at 1.65 Å resolution unveils a typical cupredoxin fold with extended loops, never observed in previously characterized cupredoxins, that might be involved in the interaction of AcoP with its physiological partners. Moreover, the structure shows that the green color of AcoP cannot be attributed to nonclassical copper ligands, its green-colored copper center raising from a long Cu-S (Cys) bond, determined by both X-ray diffraction and EXAFS. The crystal structures of two AcoP mutants confirm that the active center of AcoP is highly constrained. Comparative analysis with other cupredoxins of known structures, suggests that in AcoP the second coordination sphere might be an important determinant of active center rigidity due to the presence of an extensive hydrogen bond network.

## INTRODUCTION

Cupredoxins belong to a widely occurring family of copper-binding proteins involved in key biological processes, such as respiration, photosynthesis, the nitrogen cycle or copper homeostasis ^1,2^. They share a typical fold with a beta-sandwich involving 7-8 strands arranged into a Greek-key beta barrel. Depending on a slightly different arrangement of the N-terminal beta strands, two distinct sub-folds have been identified: the plastocyanin and the rusticyanin-stellacyanin subfamilies ^3^. The overall cupredoxin fold has been conserved throughout evolution ^3^ and can be found in all domains of life. Well known proteins containing this fold are single domain proteins, such as Type 1 (T1)- copper proteins (azurin, plastocyanin, rusticyanin, …) and cupredoxin-like proteins (CupA, CopC or PmoD, …). It can also be found in multi-domain enzymes such as multicopper oxidases (MCOs including laccases, bilirubin oxidases, *…*) and copper-containing nitrite reductases (NiRs). Despite sharing a common fold, cupredoxins can accommodate various types of copper centers with different properties related to specific functions such as electron transfer, copper sequestration, or catalysis ^4–6^. For this reason, cupredoxins represent an ideal model system for understanding the structure-function relationship of copper proteins. Among them, T1-copper proteins provide an attractive model system due to their apparent “simplicity” and therefore have been the subject of intensive studies. Indeed, T1-copper proteins are single domain cupredoxins that bind one copper atom, typically coordinated by three strong equatorial ligands which are highly conserved including one cysteine and two histidines as well as a weak axial ligand, most commonly a methionine residue^5^. One fascinating feature of T1-copper proteins is their color heterogeneity from light blue to red which results from the various geometries adopted by the copper atom and its ligands in the metal center. This gives rise to unique electronic and spectroscopic features in their oxidized form. Based on these features, T1 proteins are divided into several sub-families: blue or “classical T1” (tetrahedral), green or “T1.5” (distorted tetragonal), and red or “T2-like” (tetragonal)^7,8^.

From detailed spectroscopic analysis, combined with structural data and theoretical calculations performed on natural or “engineered” T1-copper proteins, a model which rationalizes the variation of spectroscopic properties in different sub-families has been proposed ^7^. According to the coupled distortion model, blue copper centers (T1) have a tetrahedral geometry due to a short Cu^2+^-S(Cys) bond length (~ 2.08Å), while the Cu^2+^-S(Met) bond length is around 3 Å. This leads to an intense S(Cys) π → Cu ligand-to-metal charge transfer (LMCT) transition ^2,8^ which gives rise to a strong absorption band at 600 nm. By contrast, green copper centers (T1.5) exhibit a distorted tetragonal geometry with a longer Cu^2+^-S(Cys) bond length (~ 2.22 Å) and a shorter Cu-S^2+^(Met) bond length (~ 2.5 Å). These differences are illustrated by a weaker absorption band at 600 nm as well as an additional band near 450 nm attributed to S(Cys) σ → Cu LMCT, and in some cases to a S(Met) → Cu LMCT transition. This is concomitant with a rotation of the (Cys)S-Cu-S(Met) plane with respect to the (His)N-Cu-N(His) plane ^8^. Another sub-class of cupredoxins, called “blue-perturbed” (such as rusticyanin, stellacyanin, or cucumber basic protein as CBP, *etc‥*.), has intermediate features between blue and green copper proteins, resulting from a tetrahedral distorted copper center geometry, with a Cu^2+^-S(Cys) bond length ~2.17Å ^9^ and a Cu^2+^-S (Met) bond length around 2.8 Å. As of today, only one example of T2-like cupredoxin protein (or red), called nitrosocyanin, has been found in nature, isolated and its structure solved ^10^. In this case, a histidine is the fourth ligand and a glutamate replaces one of the equatorial histidines. This rearrangement of residues results in a rotation of the Cu-S(Cys) plane and the binding of a water molecule in the equatorial plane.

Strikingly, the redox potential (Em) of T1-copper proteins and cupredoxin domains greatly vary (from + 184 mV to + 680 mV vs. SHE). Several studies based on protein engineering/site-directed mutagenesis attempted to rationalize the electronic and redox features of T1-copper proteins. From these studies, the role of the first and second coordination spheres in controlling the redox potential of T1 copper proteins was highlighted and confirmed by Lu *et al*. ^11^. Numerous studies have identified key factors regulating the electronic and redox properties of T1-copper center. Despite this, a rational link between the structure of the active site, the redox potential of the protein and spectral properties remains to be elucidated. Moreover, new proteins harboring a cupredoxin fold with intriguing features are constantly being discovered ^12–16^, illustrating broad versatility among this class of proteins.

Among novel cupredoxins, a new T1 copper protein (AcoP) from the acidophilic organism *Acidithiobacillus ferrooxidans* was isolated ^17^ and characterized in detail ^18–21^. AcoP is a membrane-associated protein with a soluble periplasmic cupredoxin domain that binds one atom of copper that is exposed in the periplasm of the bacterium. This soluble domain is anchored to the membrane by one N-terminal transmembrane segment and interacts with the terminal enzyme of the respiratory chain: the cytochrome *c* oxidase (C*c*O) ^18,22^. While the exact physiological role of this protein as an actor within the electron transfer chain is still obscure, it seemed to play an important role in maintaining the C*c*O activity under extreme acidic conditions in the periplasm (estimated to be around pH 3)^18^. Moreover, we recently demonstrated intermolecular electron transfer between the high potential heme of a dihemic cytochrome *c* (Cyt *c*) and AcoP ^20^ as well as the existence of an alternative electron transfer pathway from the Cyt *c* to C*c*O through AcoP^22^. Furthermore, AcoP’s spectroscopic properties are reminiscent of a T1.5 copper center. Based on mutagenesis studies, we demonstrated that AcoP binds one copper atom coordinated by the classic Cys-Met-His_2_ ligand coordination ^19,21^. Strikingly, AcoP exhibits an unexpectedly high redox potential (+ 566 mV at pH 5 vs SHE) ^19^ for this type of copper protein. Indeed, examples of T1.5 copper proteins found in nature are rare, such as Nirs or auracyanin D, and are usually associated with “low” redox potential (+ 250 mV and + 90 mV vs SHE, respectively) ^23,24^. Based on several mutagenesis studies, it was proposed that shortening the Cu^2+^-S(Met) bond could result in the stabilization of the Cu^2+^ state, thus lowering the redox potential ^24–26^. Other studies, made on high redox potential blue perturbed cupredoxins (*i.e*. rusticyanin), have highlighted the importance of the second coordination sphere in determining the redox potential of the protein ^27^.

In this study, we focused on structural characterization of wild type AcoP and two mutants, using X-ray crystallography, SAXS and EXAFS. Our work unveils very unusual features of the AcoP copper center geometry, reminiscent of both blue and green copper centers. The role of such features and of the first and second coordination spheres were investigated and the implications for electronic and redox properties discussed.

## MATERIALS AND METHODS

### 1.1. Protein mutagenesis, protein purification and spectroscopic measurements

Proteins expression and purification were performed as described previously ^19,21^. UV-Vis ABS spectra of purified proteins were recorded using a Cary 50 Bio (Varian) spectrophotometer. Purity and proteins concentration were determined with a theoretical molar extinction coefficient (ɛ_280_) of 25,440 M^−1^ cm^−1^.

### 1.2. Crystallization of AcoP wild-type and mutants

Crystals of wild-type (WT) AcoP, as well as of the M171A and H166A mutants, were grown using the sitting-drop vapour diffusion technique in MRC 96-well crystallization plates (Swissci). 0.1 nL of protein and precipitating solution were dispensed using a Mosquito robot (TTP Labtech) and equilibrated against 50 μL of precipitating solution. Early crystal hits were obtained starting from commercial crystallization solution kits (Molecular Dimensions Limited). In final crystallization experiments, the protein concentration was set to 8-10 mg/mL. In the case of AcoP WT and M171A, the crystallization solution contained 100 mM potassium acetate, 10 mM potassium chloride, 50 mM buffer (MES, Hepes, or TRIS, pH 6.0 to 8.0) and either PEG 3000 (34% to 44% w/v) or PEG 3350 (29% to 39% w/v) as a precipitating agent. In the case of AcoP H166A, crystals were obtained in 0.1M Bis-Tris pH 5.5 and 2.0 M ammonium sulfate. Plates were incubated at 20°C in a Rock Imager station (Formulatrix), which also allowed monitoring of crystal growth during time. Well-defined rod-shaped crystals grew for AcoP WT and M171A, with dimensions of up to 50 μm thickness and 200 μm length. These crystals were stable and diffracting even after several weeks. One year after they appeared, crystals of AcoP WT and M171A were still transparent and green respectively. This is in agreement with the high redox potential measured for AcoP WT (which tends to remain reduced even in aerobic conditions) and with our inability to reduce the M171A AcoP mutant. Small, 30 μm transparent pyramidal crystals were obtained for AcoP H166A. The crystals were cryoprotected with ~20% v/v glycerol added to the crystallization conditions. PEG cryoprotectant solutions were prepared using 150 mM sodium acetate pH 3.6 as a buffer; the final pH was however measured to be 4.6. The structures of reduced and oxidized WT AcoP were obtained from crystals soaked respectively for 75 and 120 minutes in 2 μL of cryoprotectant containing 10 mM of either sodium ascorbate or iridium(IV) sodium hexachloride and flash-cooled in liquid nitrogen.

### 1.3. Diffraction data collection and processing

X-ray data collection was performed at the ESRF (Grenoble, FR; beamlines id29, id23-1 and BM30A) and Soleil (Gif-sur-Yvette, FR; beamline Proxima1) synchrotrons. Cryo-cooled crystals were mounted under a stream of gaseous nitrogen at 100°K. All diffraction data were indexed and integrated using XDS ^28^. Subsequent data processing and interpretation relied on programs available in the CCP4 Suite (v 6.4.0) ^29^. Space group determination and scaling was achieved using Pointless (v 1.8.12) and Scala (v 3.3.21) ^30,31^. The crystal of all AcoP variants (reduced, oxidized, M171A and H166A) belong to space group P 4_1_ 2_1_ 2. A data set collected above the Cu K-edge (1.07137Å) was used for SAD phasing (single-wavelength anomalous dispersion) of the reduced AcoP. Copper sites location and refinement, density modification, and preliminary model building was obtained using the CRANK suite (v 1.5.46) ^32^. Several secondary structure elements were unambiguously identified, pruned after visual inspection, and used as a poly-alanine model for phasing a native dataset at higher resolution by molecular replacement in PHASER (v 2.5.5) ^33^. Starting from this solution, 95% of the model was assigned by automated model building in ARP/wARP (v 7.4) ^34^. The structure of the other AcoP variants (oxidized, M171A and H166A) were solved by molecular replacement with PHASER using the reduced variant as starting model. Model refinement was carried out in Refmac (v 5.8.0073) ^35^, using isotropic B factors to model atomic displacement and disorder. Anisotropic B factors and no restraints were used for Cu(I) and Cu(II) ions, in order to yield the most precise and unbiased atomic positions. Data collection and refinement statistics are reported in Table 1. Model geometry was validated using MolProbity ^36^. Atomic diffraction precision indicator (DPI) values were obtained using the Online DPI server ^37^ and converted to error estimates for the Cu-ligand bond distances as described in ^38^. Combined single crystal UV-Vis microspectrophotometry and X-ray data collection were carried out at ESRF beamline BM30 using the ESRF CryoBench microspectrophotometer ^39^.

**Table 1:** Data collection and refinement statistics.

| **Data collection** | **reduced** | **oxidized** | **H166A** | **M171A** |
| --- | --- | --- | --- | --- |
| Space group | P4<sub>1</sub>2<sub>1</sub>2 | P4<sub>1</sub>2<sub>1</sub>2 | P4<sub>1</sub>2<sub>1</sub>2 | P4<sub>1</sub>2<sub>1</sub>2 |
| a, b, c (Å) | 72.95, 72.95, 112.82 | 73.48, 73.48, 112.90 | 73.91, 73.91, 113.87 | 73.75, 73.75, 113.08 |
| $\alpha$, $\beta$, $\gamma$ (°) | 90, 90, 90 | 90, 90, 90 | 90, 90, 90 | 90, 90, 90 |
| Resolution (Å)* | 50-1.65 (1.74-1.65) | 40.0-1.7 (1.79-1.7) | 40.0-2.1 (2.21-2.1) | 50-1.82 (1.92-1.82) |
| Unique reflections* | 37412 (5387) | 34624 (4976) | 18289 (2652) | 28048 (4087) |
| Redundancy* | 6.9 (7.1) | 5.6 (5.5) | 7.3 (7.4) | 7.4 (7.5) |
| Completeness (%)* | 99.9 (99.9) | 99.7 (99.8) | 96.7 (98.1) | 98.3 (99.6) |
| I/$\sigma$* | 15.4 (1.9) | 6.1 (1.4) | 10.4 (2.0) | 14.6 (1.8) |
| R<sub>meas</sub> (%)* | 5.6 (111.1) | 16.9 (116.6) | 15.3 (124.6) | 8.5 (107.2) |
| CC1/2 | 0.999 (0.638) | 0.987 (0.485) | 0.996 (0.477) | 0.998 (0.616) |
| Mosaicity | 0.09 | 0.13 | 0.06 | 0.07 |
| **Refinement and model quality** |  |  |  |  |
| Resolution (Å) | 44.6-1.65 | 38.3-1.7 | 35.8/2.1 | 47.4-1.82 |
| Reflections | 35493 | 32890 | 17368 | 26633 |
| R<sub>fac</sub> / R<sub>free</sub> (%) | 16.0 / 19.0 | 23.7 / 26.0 | 19.1 / 24.2 | 16.0 / 19.8 |
| Number of atoms |  |  |  |  |
| protein / ion / ligand / water | 2242 / 10 / 18 / 173 | 2096 / 22 / 12 / 123 | 2101 / 2 / 12 / 109 | 2124 / 18 / 30 / 205 |
| B-factors (Å<sup>2</sup>) |  |  |  |  |
| protein / ion / ligand / water | 21.6 / 36.1 / 44.3 / 40.7 | 20.5 / 37.2 / 42.0 / 35.2 | 28.1 / 35.7 / 58.5 / 43.3 | 23.6 / 34.5 / 43.8 |
| Rmsd |  |  |  |  |
| bond (Å) | 0.015 | 0.013 | 0.013 | 0.013 |
| angle (°) | 1.930 | 1.689 | 1.840 | 1.681 |
| Ramachandran plot (%) |  |  |  |  |
| most favored | 95.7 | 95.3 | 94.2 | 95.0 |
| allowed regions | 3.5 | 3.5 | 4.7 | 4.2 |
| outliers | 0.8 | 1.2 | 1.1 | 0.8 |
| PDB accession code | 7Z3B | 7Z3F | 7Z3G | 7Z3I |

\* Values in parentheses are for the highest-resolution shell

### 1.4. EXAFS

EXAFS data collection and analysis: Samples were mixed with 20% (vol/vol) ethylene glycol and measured as frozen glasses at 10 K. Cu K edge (8.9 keV) extended X-ray absorption fine structure (EXAFS) and X-ray absorption near edge structure (XANES) were collected at the Stanford Synchrotron Radiation Lightsource on beamline 9-3 using a Si 220 monochromator with a φ= 90° crystal set and a Rh-coated mirror located upstream of the monochromator with a 13 keV energy cut-off to reject harmonics. Kα fluorescence was collected using a 100-element Canberra Ge array detector. A Z-1 metal oxide filter and Soller slit assembly was placed in front of the detector to attenuate the elastic scatter peak. A buffer blank was subtracted from the raw data to produce a flat pre-edge and eliminate residual Ni Kβ fluorescence of the metal oxide filter. Energy calibration was achieved by placing a Cu metal foil between the second and third ionization chamber. Data averaging, background subtraction, and normalization were performed using EXAFSPAK. The experimental energy threshold (k=0) was chosen as 8985 eV. Spectral simulation was carried out by least-squares curve fitting, using full curved wave calculations as formulated by the program EXCURVE 9.2 as previously described^40–42^.

### 1.5. Small angle X-ray Scattering (SAXS)

SAXS analysis was conducted at the SIBYLS beamline at Lawrence Berkeley National Lab’s Advanced Light Source. Data were collected both in size exclusion coupled SAXS (SEC-SAXS) mode and in high throughput SAXS (HT-SAXS) ^43^. The data from both measurements were merged to maximize the signal to noise in all regions. As the protein is 16.7kDa, the injection volume and concentration of 100μL at 6mg/mL produced noisy results in the high q (momentum transfer q = 4*π* (sin(θ/2))/λ, where θ is the scattering angle and λ is the X-ray wavelength of 1.23Å). The high q region was therefore supplemented with one HT-SAXS measurement on 30 μL at 6mg/mL. The sample was collected in a transmission geometry with a sample thickness of 1.5mm. Initial analysis was conducted with the ScÅtter program. The shape determined from the SAXS results are from GASBOR ^44^. The comparison of the measured SAXS data with the crystal structure was done using FoXS ^45^. To build out the structure to include missing loops the program BilboMD ^46^ was used.

## RESULTS and DISCUSSION

### X-ray structure of AcoP: Overall fold and structure

We solved the X-ray crystal structure of the soluble as-isolated form of AcoP at 1.65 Å resolution (PDB ID: 7Z3B) (Figure 1A). The lack of color of native AcoP crystals suggests that the protein is in its reduced form, as expected given its high reduction potential. The asymmetric unit of AcoP crystals contains two molecules (A and B) (Figure S1). The two polypeptides do not engage in extensive interactions and are unlikely to represent a biological dimer, consistent with solution data obtained by SAXS that point to a monomeric species in solution (Figure 1B). Indeed, the measured radius of gyration was 16 +/− 1Å using a Guinier analysis and 16.5Å from an analysis of the pair distribution function, are in good agreement with the crystal structure. The Porod exponent was 4.0 indicating a well folded globular protein. The radius of cross-section was 13 Å, close to the radius of gyration, further supporting a globular structure. The mass determined from SAXS was 13 +/− 5kDa. The maximum dimension was 53 Å. To further connect the SAXS results with the atomic resolution crystal structures, the shape was defined by the GASBOR ^44^ program and modelling was performed using FoXS ^45^ and BilboMD ^46^. A model of the full-length sequence was made based on the crystal structure. Missing loops were added and then allowed to sample conformations until a best fit was found with the SAXS data. An excellent fit with a χ^2^ agreement of better than 1 was found. The fit to the data, the model and its placement inside the calculated shape is shown in Figure 1B.

**Figure 1:**
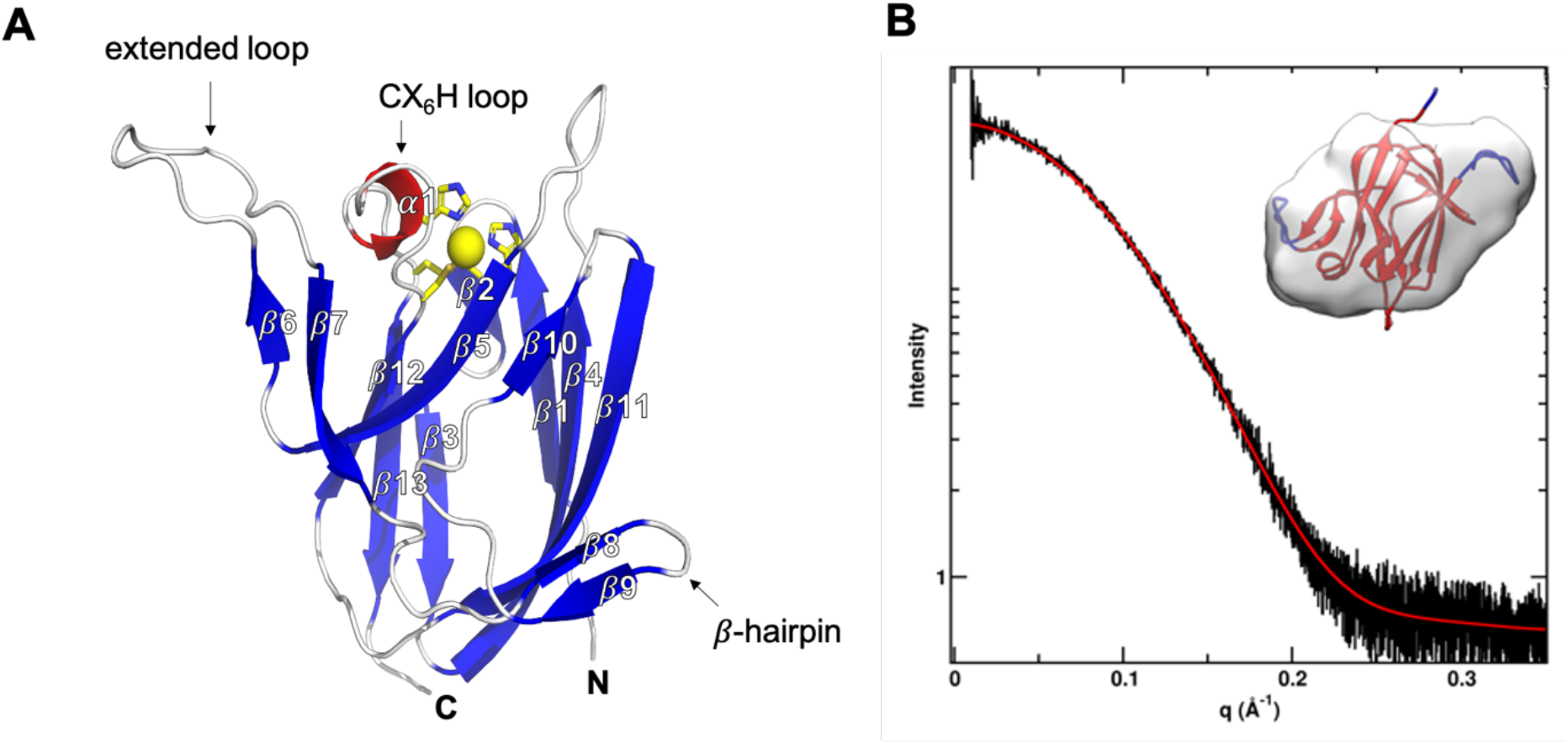
Structure and SAXS analysis of AcoP. (A) Chain A of the soluble form of AcoP in its reduced Cu(I) state. Beta strands are in blue, alpha helix in red and the copper atom in yellow, (B) SAXS data for AcoP. Reciprocal space experimental SAXS curve (black) is overlaid with the predicted scattering (red). Inset: Ab initio shape reconstruction of AcoP based on the SAXS data and overlaid with the crystal structure.

As expected, AcoP displays typical beta-barrel topology of the cupredoxin fold, with three clear differences compared to canonical structures: (i) an alpha helical loop connecting three copper ligands of an unusual length of 6 residues between cysteine and histidine (CX_6_H); (ii) on the opposite side, a small β-hairpin (res. 119-128) that extends one of the 2 beta-sheets; and (iii) an extended loop (β6-β7 loop, res. 90-114) on the same side of the CX_6_H loop (CX_6_H loop) (Figure 1).

The fold of AcoP is the same as found in the rusticyanin-stellacyanin subfamily ^3^. Structural comparison of AcoP with other well-known cupredoxins highlights the presence of a well-conserved region, corresponding to the beta sandwich of the cupredoxin fold (Figure 2 A, grey) and of a non-conserved region (residues 90-128 in AcoP) that includes the extended loop and the β-hairpins mentioned above (Figure 2 A, green). By contrast, this region is highly conserved in AcoP homologues ^19^. Hence, it can be speculated that this region might be important for specific functions or interactions with partner(s). This region is a source of molecular diversity within the cupredoxin family, as shown for Azurin, Pseudoazurin and Cucumber Basic Protein (Figure 2A). Strikingly, AcoP extended loop is reminiscent of protruding motifs observed in copper-loading chaperones which do not possess the cupredoxin fold, such as in the Sco and PCuAC protein families (Figure 2B).

**Figure 2:**
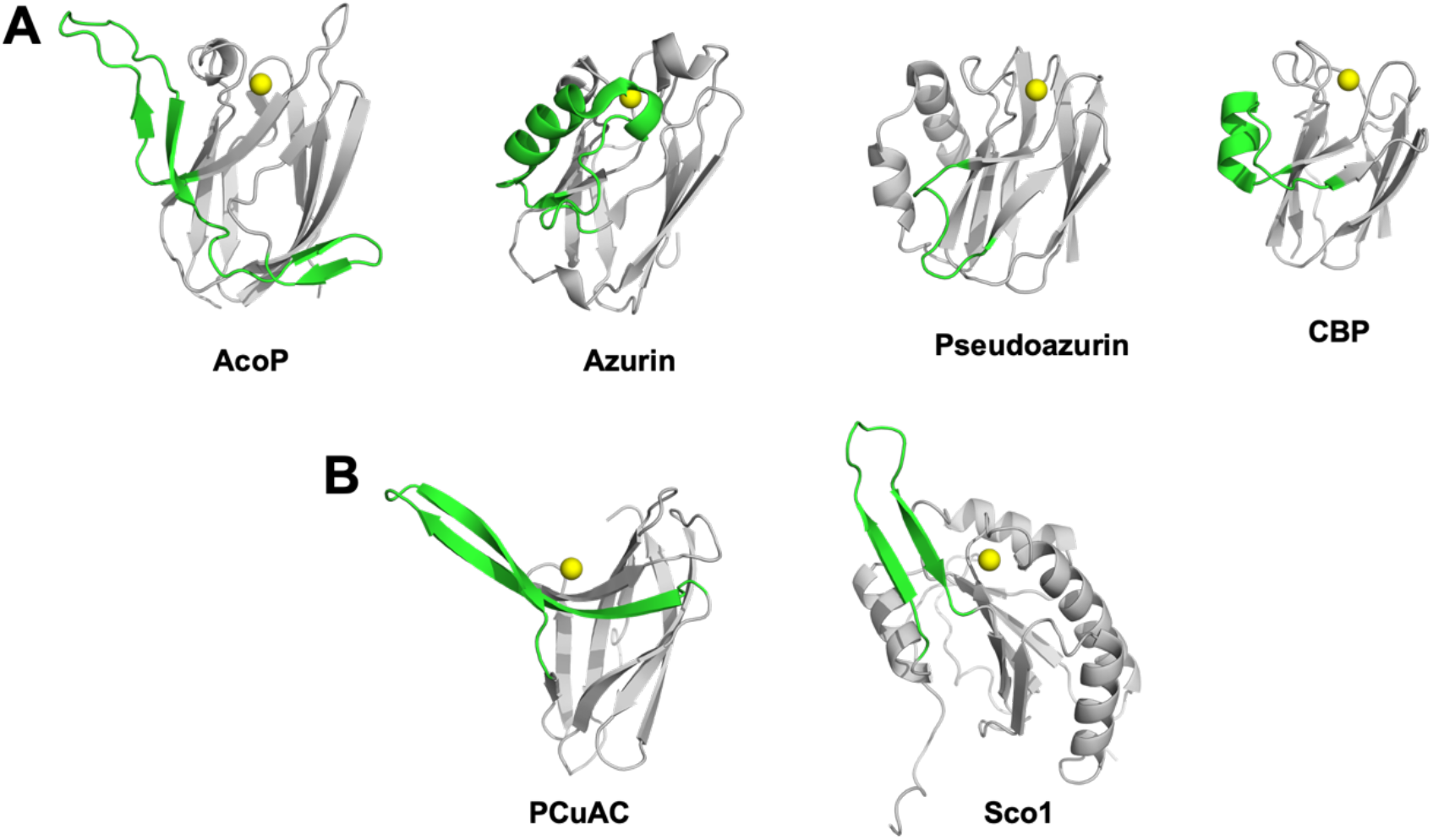
Comparaison of the AcoP fold with that of other cupredoxins and copper-loading chaperones. A) Variation of a region colored in green (residues 90-128 in AcoP) among the superfamily of cupredoxins, AcoP, Azurin (PDB 1E5Y), Pseudoazurin (PDB 1PZA) or CBP (PDB 2CBP), B) Comparison of this cupredoxin non-conserved region (colored in green) found in AcoP, with regions found in copper-bound metallochaperone such as PCuAC from *Thermus thermophilus* (PDB 2K70), and human Sco1 from PDB 2GQM.

### Examining the first coordination sphere

The crystal structure of the protein confirmed the identity of the copper ligands previously proposed based on bioinformatics analysis and mutagenesis studies ^21^. AcoP is, therefore, the first single domain green type cupredoxin with a set of classical ligands set, to be structurally characterized. The Cu-ligand distances in the reduced state determined by X-ray crystallography for each ligand are reported in Table 2/figure 3A.

**Table 2:** AcoP Cu-ligand bond distances from crystallographic and EXAFS data.

|  | AcoP red | AcoP ox | M171A ox | H166A red |
| --- | --- | --- | --- | --- |
| Cu-N(His85) (Å) | 2.00 ± 0.005<sup>+</sup><br>2.00 (0.011)* | 2.04 ± 0.01<sup>+</sup> | 2.03 ± 0.015<sup>+</sup> | 1.92 ± 0.03<sup>+</sup> |
| Cu-S(Cys159) (Å) | 2.23 ± 0.005<sup>+</sup><br>2.20 (0.009)* | 2.27 ± 0.06<sup>+</sup> | 2.25 ± 0.04<sup>+</sup> | 2.15 ± 0.015<sup>+</sup> |
| Cu-N(His166) (Å) | 2.10 ± 0.005<sup>+</sup><br>2.00 (0.011)* | 2.18 ± 0.01<sup>+</sup> | 2.05 ± 0.005<sup>+</sup> | - |
| Cu-S(Met171) (Å) | 2.92 ± 0.06<sup>+</sup><br>2.73 (0.059)* | 2.77 ± 0.03<sup>+</sup> | - | 2.80 ± 0.015<sup>+</sup> |

<sup>+</sup> Determined by X-ray crystallography (molecule A and B averaged).
\* Determined by EXAFS. Estimated values after fitting the data are listed with Debye Waller terms (2σ<sub>2</sub>) in parentheses.

**Figure 3:**
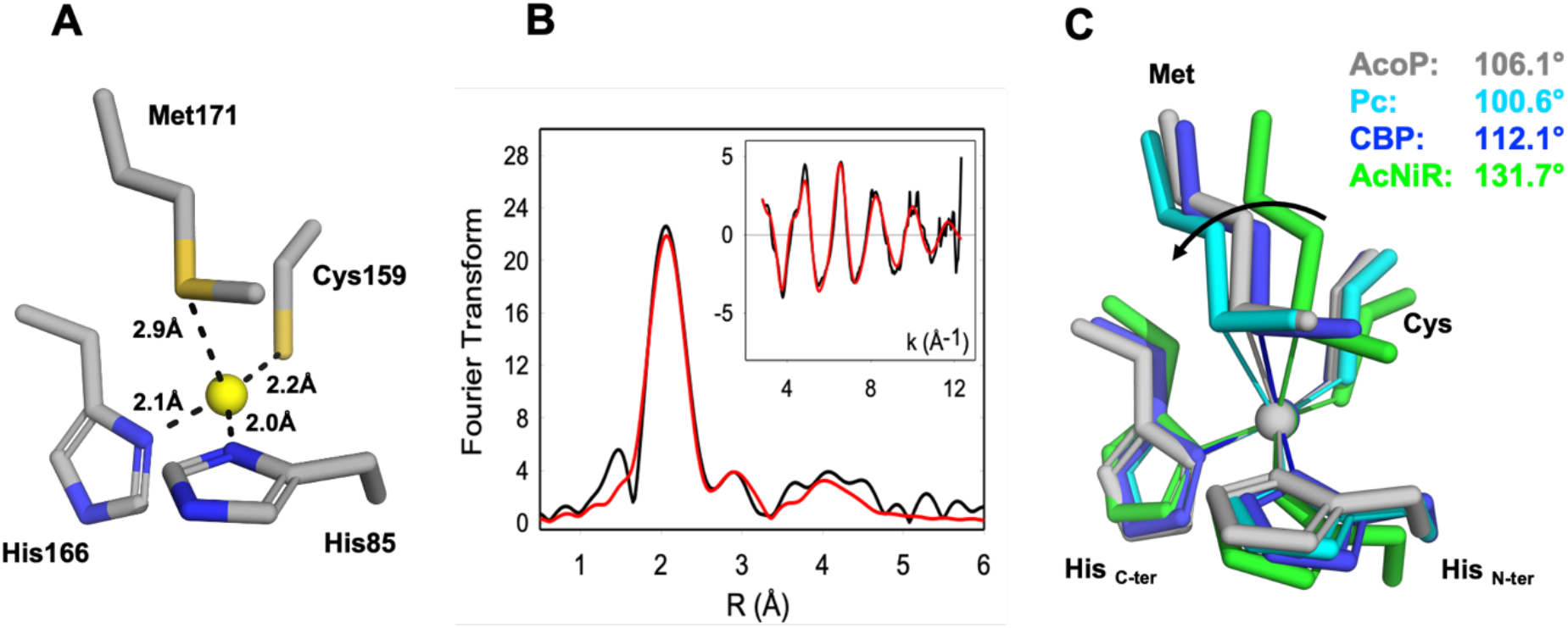
Copper center geometry of AcoP and comparison with other copper centers. (A) AcoP’s copper site, geometry and distance between Cu and the ligands in the reduced state, (B) EXAFS spectroscopy of AcoP. Fourier transform and EXAFS (inset) for AcoP as isolated protein. Black traces are experimental data, red traces are simulated data. The best fit gave metrical details, listed in table 2. (C) Overlay of the copper sites of Plastocyanin (Pc, PDB 1PLC, light blue), cucumber basic protein (CBP, PDB 2CBP, dark blue), nitrite reductase (AcNiR, PDB 1NIF, green). The arrow indicates the variation of the angle N(His_C-terminal_)-Cu-S(Met) depending on the cupredoxin type and the values of the angles are indicated.

Bond lengths between the Cu atom and the histidine ligands are typical of T1 copper center (expected from 1.9 Å to 2.2 Å). Cu-S(Cys) distance was measured to be ~2.23 Å for AcoP. Strikingly, the distance between the Cu atom and the Met171 axial ligand is similar to the distance observed for classic blue copper sites (amicyanin ̴ 2.91 Å (PDB 2RAC), plastocyanin ̴ 2.87 Å (PDB 5PCY), pseudoazurin ̴ 2.7 Å (PDB 8PAZ) ^1^. A distance of 2.43-2.55 Å was previously reported for the green copper center of CuNiR ^47–49^. Those distances have been determined on the AcoP reduced form, once oxidized, small rearrangements in the copper center geometry could occur even though cupredoxins are well known to be rigid and maintain similar geometry between reduced and oxidized species in order to facilitate electron transfer via a small reorganisation energy ^50^. To test if distances between the Cu atom and its ligands vary depending on AcoP redox state, we soaked the crystal into an oxidizing agent, a green color crystal was obtained indicating the oxidation of at least some of the protein molecules within the AcoP crystals to the Cu^2+^ form. As photoreduction of metallocenters by exposure to X-rays (*via* the generation of solvated photoelectrons) has been well described ^51^, we collected X-ray crystallographic data and performed online crystal microspectrophotometry to monitor such reduction, Cu^2+^ AcoP crystals showed spectroscopic features which were equivalent to oxidized AcoP in solution ^19^ with absorption maxima at 438 and 568 nm (Figure S2). The X-ray doses associated with a typical crystallographic data collection would likely lead to a fully reduced crystal, and this photoreduction phenomenon was confirmed on AcoP (Figure S2A). We consequently decided to use a helical data collection strategy to minimize photoreduction. Crystal spectra were measured before and after helical data collection producing spectra that were superimposable (Figure S2B). This result validates that the structure solved with helical collected data corresponds to the oxidized form of AcoP. After refinement, no difference between the structures of the reduced and partially oxidized forms of AcoP could be observed (rmsd ~0.36 Å), suggesting that as expected for cupredoxins no or only a very small modification of the distance between copper and its ligands occur and the distance between Cu-S(Met) remains relatively long for a green type copper center.

The value of the Cu-S(Cys) distance has been interpreted as a strong predictor of the sub-class of cupredoxin since it is dependent on the degree of covalency of the Cu-S(Cys) bond ^40,52^. Blue copper proteins such as plastocyanin and azurin exhibit short Cu-S(Cys) distances (2.07 – 2.15 Å) as the result of strong covalency associated with the d_x^2^_-_y^2^_ to S-p π-bond which is largely responsible for the low energy ~600 nm S(π) to Cu(II) charge transfer band with a weaker S(σ) to Cu(II) at higher energy. Red copper proteins exemplified by nitrosocyanin and Sco exhibit normal Cu-S distances around 2.25 Å and dominant higher energy LMCT bands between 350 and 400 nm due to ligand-σ to Cu(II) charge transfer. Green copper proteins such as nitrite reductase have intermediate LMCT behavior where weakened S-p π together with lower energy ligand-σ interactions result in two bands of almost equal intensity between 460 and 500 nm. Cu-S(Cys) distances for green cupredoxins are typically in the range 2.18 – 2.22 Å nojiri ^49^. The ~2.23 Å Cu-S distance measured in the crystal structure fits with the distance of a green type copper center. However, the small difference between green and blue copper center in term of Cu-S(Cys) distance cannot be resolved at the resolution of the structures presented here as it falls within the uncertainty in bond length. To get precise values of the Cu-S distances, EXAFS studies were performed on the AcoP as isolated and on the reduced protein and the results are shown in Figure 3B. The data were fit to a chemical model based on the blue copper binding motif, namely a His + Cys coordination environment with potential additional coordination by methionine thioether. For the as-isolated protein best fits were obtained with two histidine residues Cu-N(imid) = 2.0 Å, and one cysteine residue with Cu-S(Cys) = 2.20 Å. Inclusion of a Cu-S(Met) interaction at 2.73 Å improved the quality of the fit by ~10 percent. However, given the large body’s of literature on EXAFS of cupredoxins where Cu-Met interactions are not observed, coordination of a Met residue at this distance must be viewed with caution, particularly as the Debye-Waller (DW) factor for the methionine shell is large. We also analyzed the data for the presence of a Cu-Met interaction at shorter distances, but the Cu-S distance always refined back to 2.7 Å with a high DW factor. Thus, we may conclude that the EXAFS data support a cupredoxin-like Cu center with the usual caveat that the Cu-S(Met) distance is poorly defined. Simulations of the reduced AcoP give almost identical metrical parameters. Therefore, the Cu-S distance of 2.20 Å found by EXAFS for AcoP is consistent with the “green copper” classification. However, both techniques, crystallography and EXAFS, agree on a Cu-S(Met) distance greater than 2.7 Å. This long Cu-S(Met) distance is unexpected for a green type cupredoxin. Indeed, several studies based on natural or engineered T1 copper centers have helped to rationalize the changes occurring in T1 copper center from blue, green and red copper sites^25,53,54^, giving rise to the coupled distortion model, which describes the interplay between the strength of the copper-axial ligand interaction and the copper-Cys covalency ^48^. According to this model, stretching the Cu-S(Cys) bond while shortening the Cu-axial ligand bond is usually accompanied by the rotation of the (Cys)S-Cu-S(Met) plane with respect to the (His)N-Cu-N(His) plane resulting in a distorted tetragonal geometry. The extent of such distortion can be quantified by measuring the N(His)-Cu-S(Met) angle ^55^. Hence, blue copper sites with tetrahedral to slightly distorted tetrahedral geometry (such as the plastocyanin and cucumber basic protein, respectively) exhibit smaller N(His)-Cu-S(Met) angles, than a green copper site with tetragonal geometry (such as CuNirs) (Figure 3C). Strikingly, the green copper center of AcoP exhibits a distorted tetrahedral geometry that closely resembles that of CBP, that belongs to the subfamily of blue perturbed cupredoxins (Figure 3C).

Overall, these experiments demonstrated that AcoP does not follow the coupled distortion theory. The AcoP green color is due to the Cu-S(Cys) distance, however, the Cu-S(Met) is not shortened. In comparison with other known green cupredoxins, AcoP is the first green cupredoxin that exhibits such a long distance between Cu and the axial residue.

### AcoP possesses a highly rigid copper center that is not significantly modified by mutations of ligand residues

To better understand the role played by the copper ligands on the geometry of the copper center and the electronic properties of the protein, the X-ray crystal structure of AcoP H166A and M171A variants were also solved. As we showed in a previous study, the replacement of the His166 residue located in the C-terminal loop as well as of the Met171 axial ligand by an Ala residue, does not affect copper binding nor overall fold of the protein ^21^. Nevertheless, this resulted in drastic modification of the spectroscopic and redox properties of the proteins ^21^. Indeed, the mutation of the His166 to Ala turned the protein into a Cu^+^ copper binding protein, while the replacement of the Met171 residue by Ala residue turned the protein into a Cu^2+^ copper binding proteins with ‘red’ spectroscopic properties ^21^. Although this early work emphasized that copper ligands indeed played a crucial role in determining the electronic and redox properties of AcoP, the molecular determinants for such features remained unexplored. Here, we described their crystal structures and discuss the structural origins of their properties. First, the structure of the His166Ala variant was solved at 2.1 Å resolution. The tertiary structure was essentially unchanged from that of native AcoP (rmsd ~ 0.45 Ǻ). The copper atom is coordinated by only three ligands in a distorted trigonal geometry with the closest atom of Ala166 being some ~3.5 Ǻ away from the Cu (Figure 4). This result confirms our previous assumption based on the characterization of this mutant, which gives rise to a silent form in UV-Vis spectroscopy due to the stabilization of the AcoP reduced form in a tri-coordinated fashion ^21^. This result is consistent with previous mutational studies of histidine ligands in cupredoxins (*e.g* H145A Nir, H117A Az et H85A Rus). In the case of the Met171Ala variant, the X-ray structure for this variant was obtained at 1.82 Å resolution. As expected, no change of the overall structure was observed (rmsd ~ 0.38 Å). In this case however, the copper center is penta-coordinated with the Met171 axial ligand replaced by an acetate molecule whose both oxygen atoms can interact with copper. The presence of this acetate is not a crystallographic artefact as it is consistent with our previous spectroscopic results where we observed a red copper center which can be modified by protein acidification resulting most likely in the removal of the acetate molecule ^21^. In both AcoP mutants, the replacement of one residue has no impact on the three others ligands since no change between the Cu atom and the other ligands was observed in both variants. Moreover, the distance between the most distant oxygen atom from the acetate molecule and the copper (Cu-O(Ace) = 2.96 Å) is also similar to the distance between the sulfur atom (from the methionine) and the copper in the wild type form Cu-S(Met) = 2.92 Å. Therefore, the structural comparison between the wild type AcoP and the mutants suggests that the copper center is relatively rigid as mutation of one residue from the first coordination sphere has limited influence on its geometry. To explain such behavior, we propose that the second coordination sphere of the copper center might be involved in the stabilization of the first coordination sphere, thus participating to the rigidity of the copper center.

**Figure 4:**
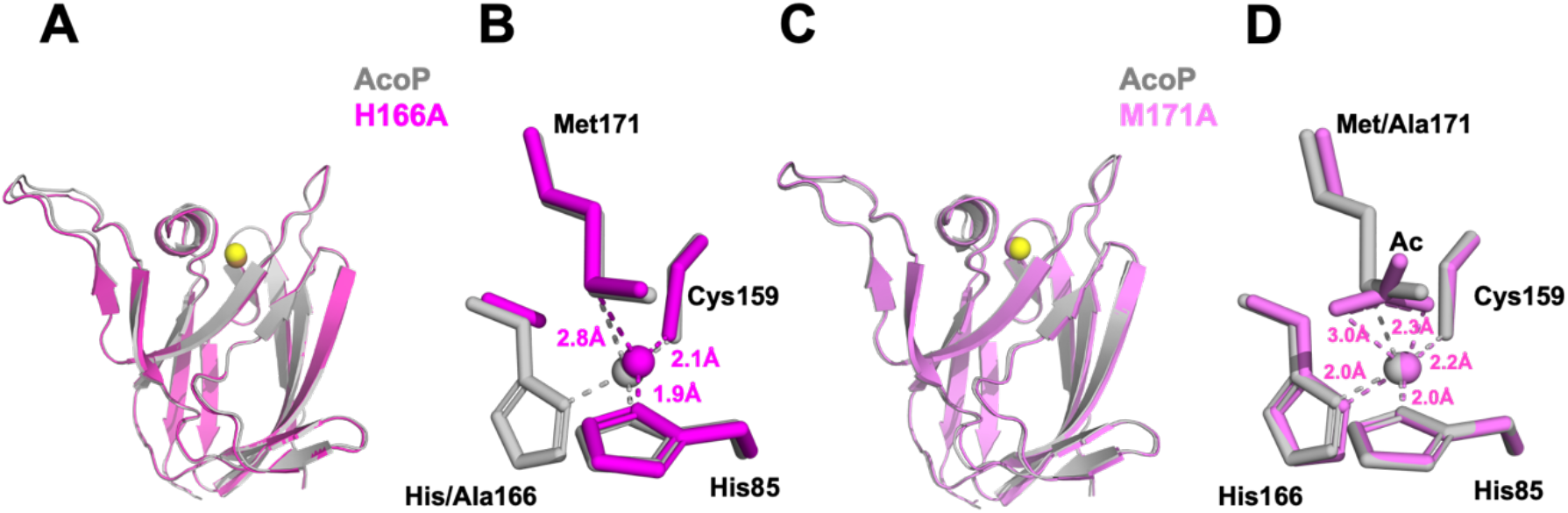
Structure of AcoP’s mutants. (A) Structure of AcoP His166Ala mutant (in purple) in comparison with wt AcoP (in grey) and superposition of their copper center (B), (C) Structure of AcoP Met171Ala mutant (in pink) in comparison with wt AcoP (in grey) and superposition of their copper center (D).

### The second coordination sphere of AcoP is designed to constrain the copper center

As shown in the previous section, the mutation of two AcoP copper ligands does not significantly affect distances and geometries between copper and the remaining ligands. This is a direct observation of the extreme rigidity of the AcoP metal center, that we previously hypothesized based on spectroscopic data ^19,21^. Another feature of AcoP is its unusually high redox potential, the highest reported to date for a green copper site ^19^. Changes in the first coordination sphere are known to influence redox potential. However, we confirm in this study that this is not the case, since AcoP has a “classic” set of copper ligands. This is also true for rusticyanin, whose high redox potential has been attributed to extensive hydrogen bond networks and to the packing of hydrophobic residues in the vicinity of the copper binding site ^27^. Mutations in the second coordination sphere have also been reported to increase cupredoxin redox potential up to a value of 1V^11^. In order to understand the structural determinants responsible for the rigidity of the copper center in AcoP, we decided to analyze the AcoP three-dimensional structure beyond the first copper coordination sphere. Starting from each copper ligand, we looked for hydrogen bond-connected neighbors within the second coordination sphere and continued searching two more levels of residues connected to the copper ligands within a continuous hydrogen bond network. We then compared the network with the ones found in Rusticyanin, Azurin and Pseudoazurin (Figure 5). It can be easily seen that in AcoP, the C-terminal copper ligands (especially histidine and cysteine) are stabilized by a large network involving 9 hydrogen bonds and 7 interconnected partners, against 7 and 6 for Rusticyanin, and 5 and 4 for Azurin and Pseudoazurin. Stabilization through hydrogen bond networks of the N-terminal copper ligand (histidine) also differs among proteins. An intra-domain hydrogen bond network, practically absent in Azurin and Pseudoazurin, relies on 8 hydrogen bonds and 9 connected partners in AcoP (only 3 bonds and 4 partners in Rusticyanin). In AcoP two other domains, regions 51-53 and 134-140 further contribute 6 and 4 hydrogen bonds, respectively, to consolidate the hydrogen bond network around His 85. In general, compared to the other three cupredoxins, the second coordination sphere of AcoP involves a larger number of residues that are two or three hydrogen bonds away (orange and red in Figure 5) from copper ligands. Further on, these residues are highly interconnected, as evidenced by the N-terminal domain, whose extensive hydrogen bond network is further restrained by long backbone connectivity (residues 74 to 85) and by multiple connections to domains 51-53 and 134-140. In view of these results, the AcoP second coordination sphere seems to provide two unexpectedly extended hydrogen bond networks, that might be a key for the exceptional rigidity of AcoP copper center.

**Figure 5:**
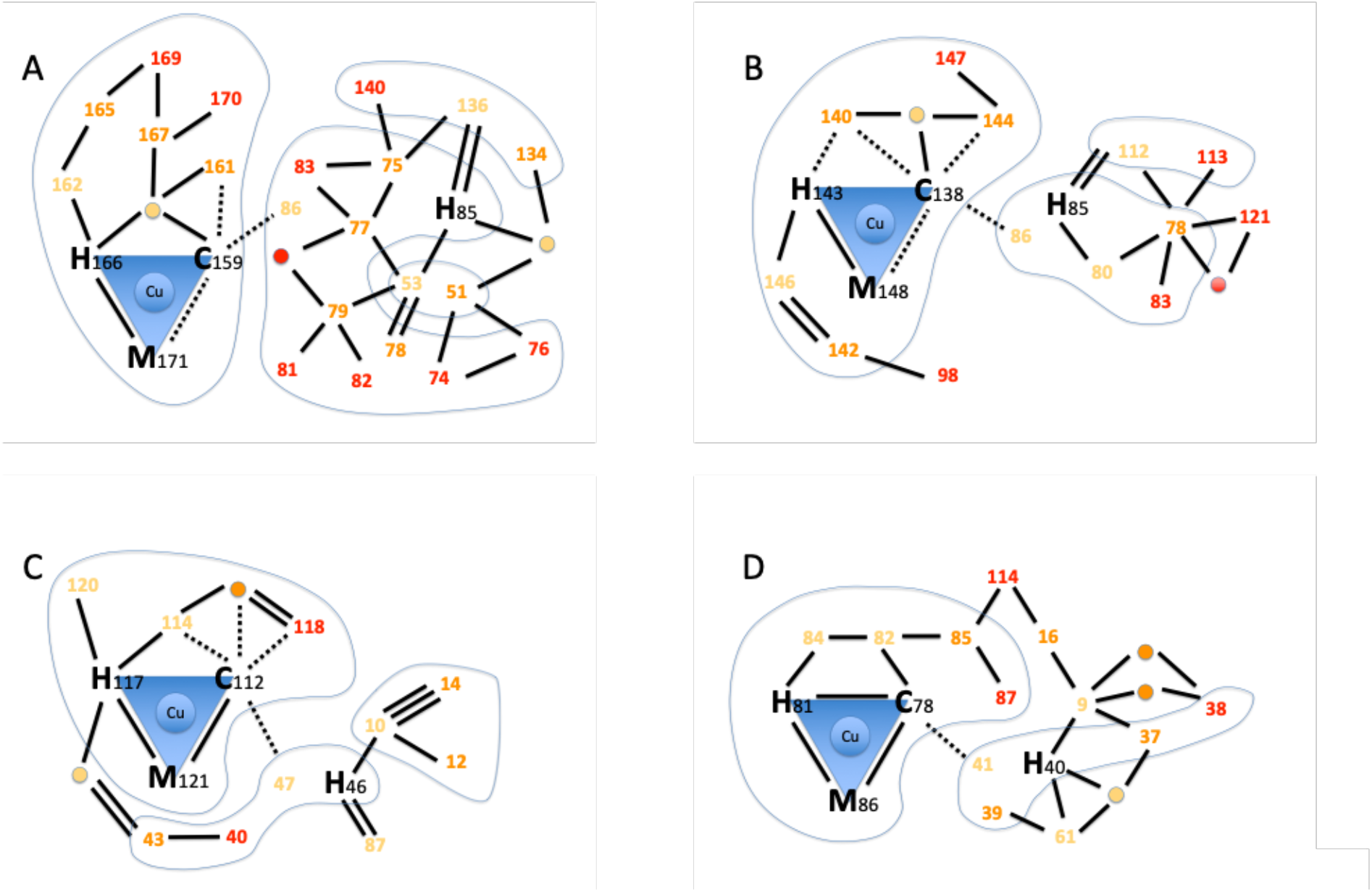
Hydrogen bond networks in cupredoxins. The scheme highlights the intricate hydrogen bond networks involving the first and second coordination spheres. Amino acids within the first coordination sphere (copper ligands) are represented by their one-letter code followed by the sequence position number. Amino acids within the second coordination sphere are represented only by the sequence position number, colored in yellow, orange or red to represent further distance (degree of indirect hydrogen bonding) to copper ligands. The same color scheme is used for bridging water molecules that participate to the network (yellow, orange or red circles). Blues lines encompass domains connected through backbone. No further search was pursued on residues (red) that are three hydrogen bonds away from copper ligands. Hydrogen bonds (2.5-3.2 Å, black lines) were used to identify network amino acids. For the first coordination sphere only, also weak hydrogen bonds (3.2-4.4 Å) are traced (dashed lines). No further search was pursued on residues connected through weak hydrogen bonds only. Blue shapes group together residues that are connected through backbone. A) AcoP (PDB: 7Z3B). B) Rusticyanin (PDB: 2CAK). C) Azurin (PDB: 4AZU). D) Pseudoazurin (PDB: 8PAZ).

## CONCLUSION

In this study, we solved and analyzed the structures of wild-type AcoP and two copper-ligand mutants. Structural comparison with other cupredoxins highlights the presence of an extended loop, previously detected by sequence alignment ^19^, which appears to be conserved in related cupredoxins from acidophilic bacteria. This loop could be important for the functional role of AcoP which is not yet fully understood. Indeed, *in vitro* studies suggested that AcoP could be an electron donor of cytochrome *c* oxidase (C*c*O) and/or an accessory protein, with chaperone-like properties, capable of preserving the integrity and activity of C*c*O metal centers in acidic environments ^18,20^. Based on this second role and on the presence of this additional loop, it is interesting to note that several types of structurally unrelated protruding motifs are also found in copper-loading chaperones. These extended motifs have been proposed to play a role in the unfolding of domains in the vicinity of the metal center to facilitate copper loading and/or disulfide-reduction of their partner protein ^56^. Similarly, the extended loop found in AcoP might play a role in partner maturation and/or protection (such as CoxB, the subunit II of *A. ferrooxidans* C*c*O). Intriguingly, Sco chaperones that are required for CoxB maturation in other species ^57^ are not found in the genome of *A. ferrooxidans*, raising the question of how CoxB maturation occurs in this species. Further studies are needed to experimentally validate the function of this loop in AcoP, and of analogous extensions in Sco and PcuAC.

The structure of AcoP validates the identity of its copper ligands ^21^. Unlike green type auracyanin D which possesses an axial glutamine residue ^58^, AcoP has a classical set of cupredoxin ligands, with an axial methionine ligand. The structure of AcoP provides insight into the structural determinants of the spectroscopic properties of this green copper cupredoxin. Previous spectroscopic analysis and site-directed mutagenesis suggested that AcoP might present a green T1.5 copper center ^19,20^. Cu-ligand distances determined by X-ray and EXAFS agree and show a Cu-S (Cys) distance of 2.2 Å. Such a distance can explain the green UV/Vis spectrum of AcoP, as a result of concomitant weakening and strengthening of *π* and σ interactions respectively. In contrast, the measured Cu-S (Met) distance (2.7 Å by EXAFS; 2.9 Å by X-ray) was unexpected. In fact, based on the coupled distortion model, it is predicted that, in tetracoordinate cupredoxins, loosening of the Cu-S(Cys) bond should be accompanied by shortening of the Cu-S(Met) bond, as measured in previously characterized green type cupredoxins ^8^. In contrast, distances measured for AcoP suggest that this cupredoxin does not follow the coupled distortion model as the long Cu-S(Cys) distance does not lead to a short Cu-S(Met) distance. Although this result is surprising for classical T1 and T1.5 copper centers, recent results obtained on *de novo* constructs that recreate blue, green and red copper centers brought by an alpha helicoidal fold, also do not respect the coupled distortion model ^59^. In this study, the Cu-S(Cys) distance is not shorter in blue compared to green constructs.

Recently, two more single-domain green-type cupredoxins have been characterized by spectroscopy: CopI from *Rubrivivax gelatinosus* ^16^ and the chimeric protein Az-TtCuA ^55^, also characterized by a classical set of copper ligands. Measuring of Cu-ligand distances in those two cupredoxins might help clarifying the relationship between spectroscopic properties, metal center structures and the coupled-distortion model in green-type cupredoxins.

The deviation of AcoP Cu center geometry from the coupled-distortion model could maybe explain AcoP redox potential, which is unexpectedly high for a green cupredoxin. Indeed, short Cu-S(Met) distances have been proposed to stabilize the Cu^2+^ ^24–26^ and as such lower cupredoxin redox potentials. Cu-S(Met) distance in AcoP is comparable to those found in blue cupredoxins, and it is therefore tempting to speculate that AcoP high redox potential might derive from a long Cu-S(Met) bond. Similar results were also obtained from DFT calculations, that successfully identified a few structural determinants of cupredoxin redox potentials, including the Cu-S(Met) distance ^60^. In this study, after summarizing the individual contributions of each structural determinant located within 6 Å of the Cu atoms, several redox potentials were predicted that well matched experimental ones. Knowing the AcoP structure, the DFT calculation on AcoP might be of interest.

The structures of two mutants of the first coordination sphere, AcoP H166A and AcoP M172A, were also successfully solved. These structures show very little variation of the metal center geometry compared to wild-type AcoP, even in the absence of one of the Cu ligand, and validate our previous hypothesis that AcoP has a highly constrained copper center: mutation of a ligand does not significantly affect the metal site structure, nor copper binding to the three remaing copper ligands^21^. In line with this, as we previously showed, even the addition of exogenous ligands to the M121A (axial) mutant had little effect on UV/Vis spectroscopic properties of AcoP, including ligands (such as dimethylsulfide) that could be expected to generate a blue copper center ^21^. In this previous study, we hypothesized that the protein scaffold might play a critical role for rigidifying AcoP green copper center. Herein, we performed detailed structural analysis and comparison of AcoP and a few model cupredoxins of known structure which suggest a key role of the second coordination sphere in constraining AcoP active center. A highly structured and extensive hydrogen bond network is in fact clearly observed in AcoP but not in the compared proteins. In other studies, the protein scaffold was also suggested not to be a passive entity, but rather play an important role in cupredoxin metal center geometry ^1^. AcoP perfectly illustrates this hypothesis. The role, if any, of constrained and constraining protein scaffolds in cupredoxins is, in our view, still an open question. We might wonder if highly structured and rigid polypeptide domains have an effect on cupredoxin redox potential. Our study cannot give a final answer, but the question seems relevant. Future studies, including site-directed mutagenesis, are needed. Previous experiments targeting hydrogen bonds in other cupredoxins already emphasized their key role in tuning redox potential ^11,61^. The unprecedented spectroscopic, electronic, redox and structural properties of AcoP suggest that this protein has the potential to become an interesting model system for future studies aiming at gaining new insight on structure-function relationship in cupredoxins.

## ACKNOWLEDGEMENTS

Authors want to thank the staff of the ESRF and Soleil synchrotrons for beamline allocation and for their assistance with single-crystal spectroscopy experiments.

SAXS data were collected at the SIBYLS beamline in Berkeley California supported by a National Institute of Health R01 GM137021-02 and the United States Department of Energy Biological and Environmental Research (DOE-BER) IDAT grant.

The authors also want to thank X. Wang, L. Zuily, N. Lahrach and M. Bauzan for technical support, E. Lojou, P. Dorlet from BIP-CNRS Marseille for fruitful discussions and V. Pecoraro for critical reading of the manuscript.

