## Supplemental file for "Three-dimensional structure of the single domain cupredoxin AcoP"

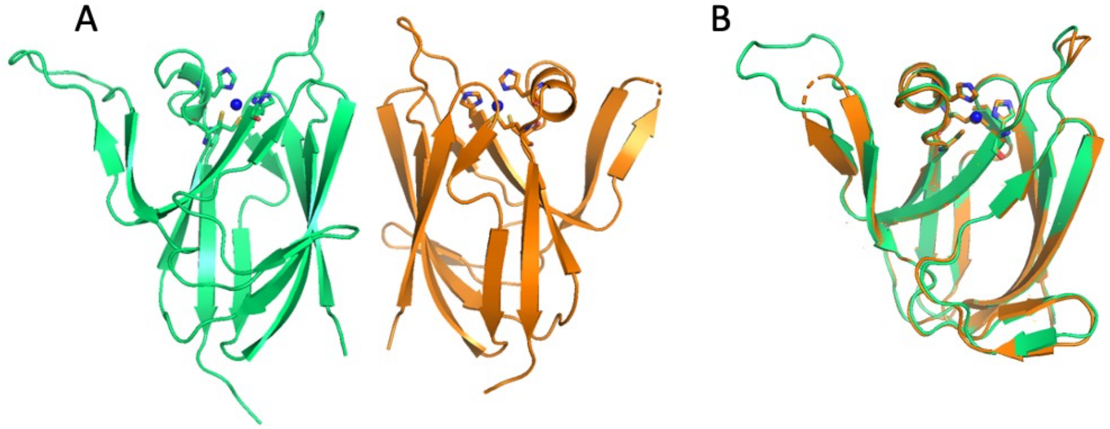


**Supp Figure 1: Structural comparison between molecule A and B of wild type AcoP.** A) The two molecules present in the asymmetric unit, and B) their superposition are displayed (molecule A in green, molecule B in orange).

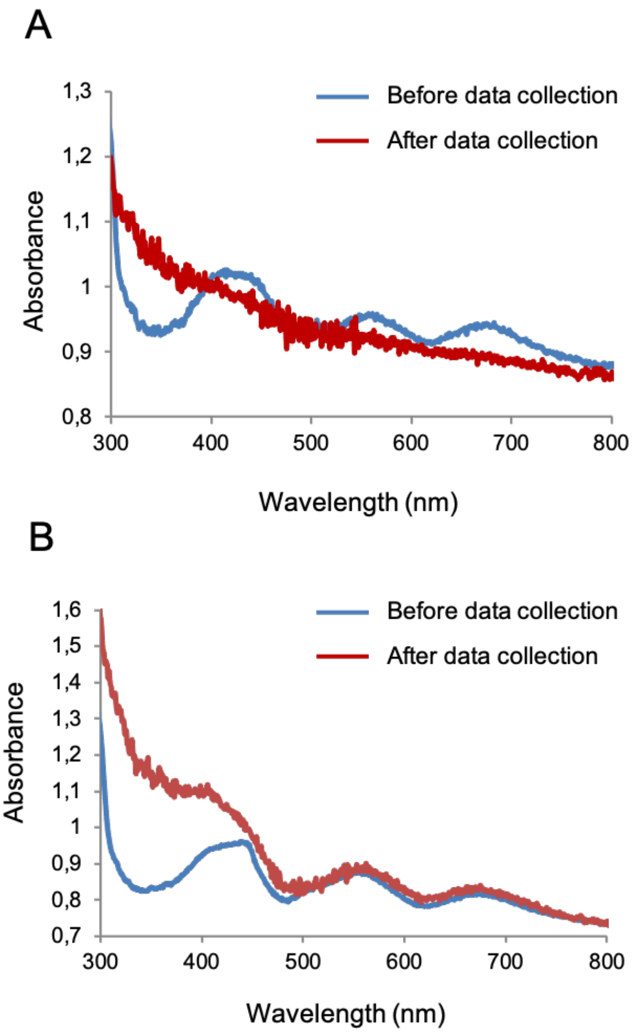


**Supp Figure 2 : Online microspectrophotometry of AcoP oxidized crystal.** (A) spectra of AcoP oxidized crystal before data collection (blue) and after data collection (red) (B) spectra of AcoP oxidized crystal before data collection (blue) and after helical data collection (red). Note an increase in absorbance at shorter wavelengths, this is a common feature in the spectra of X-ray-exposed protein crystals.
